# Automated spermatogenic staging in PAS-stained testes of Sprague-Dawley rats using a deep learning model for normal and atrophied tissues

**DOI:** 10.1101/2025.11.06.687107

**Authors:** Da-Mi Kim, Jin-Hyung Rho, So-Young Wee, Hwa-Young Son

## Abstract

The spermatogenic stage serves as a vital criterion for assessing normal spermatogenesis and is central to evaluating reproductive toxicity. Current manual methods for spermatogenic stage evaluation are time-intensive, require expert knowledge, and are less effective in detecting subtle changes or comparing stage frequencies across samples. To overcome these limitations, this study introduces a method leveraging the object detection models, Region-based Convolutional Neural Networks (R-CNN), for efficient and accurate spermatogenic stage evaluation. 14 stages were identified using Periodic Acid-Schiff (PAS)-stained Sprague-Dawley (SD) rat testicular tissue, and the approach was further applied to atrophied testicular samples as a real-world example. The model achieved a mean average precision of 0.869 and a mean average recall of 0.977 in detecting spermatogenic stages and atrophy. Agreement with pathologist assessments exceeded 91%, providing objective benchmarks for stage evaluation and facilitating the comparison of stage frequencies across multiple samples. In atrophied tissues, the model enabled quantitative grading by analyzing proportional changes in atrophied seminiferous tubules relative to normal tubules. This automated approach reduces the workload of pathologists while delivering rapid and precise assessments of toxicological changes in spermatogenesis. By integrating deep learning models, this study enhances both the accuracy and efficiency of pathological evaluations, offering a transformative tool for reproductive toxicity studies.

## Introduction

Spermatogenic staging is a critical evaluation criterion in non-clinical reproductive toxicity studies, offering an indirect assessment of whether spermatogenesis is proceeding normally. Each seminiferous tubule’s stage is classified based on its spermatogenic state, with abnormalities often reflected in changes in the frequency of specific stages. The Organization for Economic Cooperation and Development (OECD) guidelines for reproductive toxicity testing emphasize the importance of histopathological evaluation of spermatogenic stages and interstitial testicular cell structure (1, 2). However, manual evaluation of the spermatogenic stages in hundreds of seminiferous tubules per testis is highly time-intensive, requires specialized expertise, and has limited sensitivity for detecting subtle changes or comparing stage frequencies across multiple samples. Recent advancements in deep learning-based computer vision have shown significant promise in addressing such limitations. The digitization of whole slide images (WSIs) has further enhanced the applicability of deep learning models in pathology (3). In pathology, deep learning-based diagnostic tools are already being utilized to improve efficiency and accuracy in cancer diagnostics and are increasingly explored in non-clinical research (4–6). Prior studies have demonstrated the potential of deep learning models to automate spermatogenic stage evaluation of laboratory animals, reducing reliance on expert pathologists and providing precise quantitative data (7–10). Especially, Creasy et al. (2021) established the deep learning model for spermatogenic stage assessments in rats (7). Compared to earlier studies that employed separate segmentation and classification models, this study adopted a single object detection model capable of performing both localization and classification simultaneously. The algorithm implemented in this study belongs to the Region-based Convolutional Neural Network (R-CNN) family, a representative class of two-stage detectors known for their higher accuracy compared to one-stage detectors, despite their relatively slower processing speed (11). Additionally, transfer learning was performed using two widely used pretrained models from the R-CNN family—Faster R-CNN (12) and Cascade R-CNN (13)—and their performance was compared. Finally, the best-performing model was applied to WSI inference, and the results were compared with those of pathologists to assess the practical feasibility of the model in real-world scenarios.

The importance of spermatogenic staging lies in its ability to provide information about the spermatogenesis occurring within the seminiferous tubules. Spermatogenesis is the process by which spermatogonia, located on the basement membrane of seminiferous tubules, undergo mitotic and meiotic division to produce spermatozoa. Mature spermatozoa are released into lumen of seminiferous tubules (14). The spermatogenic stage of a seminiferous tubule is determined by the progression state of spermatogenesis and is classified into 14 stages in rats (15). Rats are commonly used animals in non-clinical experiments and demonstrate relatively regular sperm development, often exhibiting one predominant stage per cross-section of the seminiferous tubule, making them suitable for stage evaluation. The evaluation of 14 spermatogenic stages requires Periodic Acid-Schiff (PAS) stain, which highlights polysaccharides in acrosomes of spermatids. Traditionally, detailed spermatogenic stage assessment of seminiferous tubules has been accomplished through PAS staining (16). In contrast to previous studies using Hematoxylin and Eosin (H&E) staining, our study used PAS-stained rat testis samples to identify all 14 spermatogenic stages. Furthermore, the study explored the model’s capability to detect atrophy as an example of its real-world application. A significant pathological alteration affecting spermatogenesis is seminiferous tubule atrophy, which can be induced by various reproductive toxicants (17–19). The pathogenesis of seminiferous tubule atrophy involves damage to Sertoli cells, cytotoxicity, hypoxia, and inflammation, which can lead to germ cell depletion and, in severe cases, a reduction in testis size or weight (20). Some toxicants selectively affect specific spermatogenic stages, necessitating the evaluation of stage frequency alterations and the quantification of atrophied seminiferous tubules to assess the severity (21, 22).

This study distinguishes itself from prior research in three key aspects. First, rather than employing separate segmentation and classification models, it utilizes an object detection model from the R-CNN family, enabling simultaneous localization and classification. Second, PAS-stained rat testis tissue samples were used to assess the model’s ability to distinguish the 14 spermatogenic stages in normal testes. Third, to explore the model’s applicability in real-world scenarios, the dataset included atrophied seminiferous tubules, allowing evaluation of the model’s performance in detecting and grading pathological changes. By incorporating these elements, this study aims to advance the automated assessment of spermatogenic stages and facilitate objective, high-throughput analysis of testicular histopathology in toxicologic research.

## Materials and methods

### Image processing and annotation

#### Digital image processing

Corestemchemon Inc. provided 16 PAS-stained testis slides of 21-week-old Sprague-Dawley rats, consisting of 12 normal and 4 atrophied testes. The animals were acquired from Orient Bio Inc., and PAS stain kit was purchased from ScyTek Laboratories Inc.. Transverse sections of the right testis were used. A total of 16 slides from 16 animals were scanned at 40× magnification using a 3DHistech Ltd. slide scanner. 10 digitized whole slide images (9 normal testes and 1 atrophied testis) were cropped into 4096×4096×3 resolution tiles using the OpenSlide library (23) to facilitate deep learning model labeling. The remaining 6 whole slide images were reserved for inference.

#### Image annotation

A total of 683 tile images were annotated using Roboflow software as shown in Fig 1.

**Fig 1.**
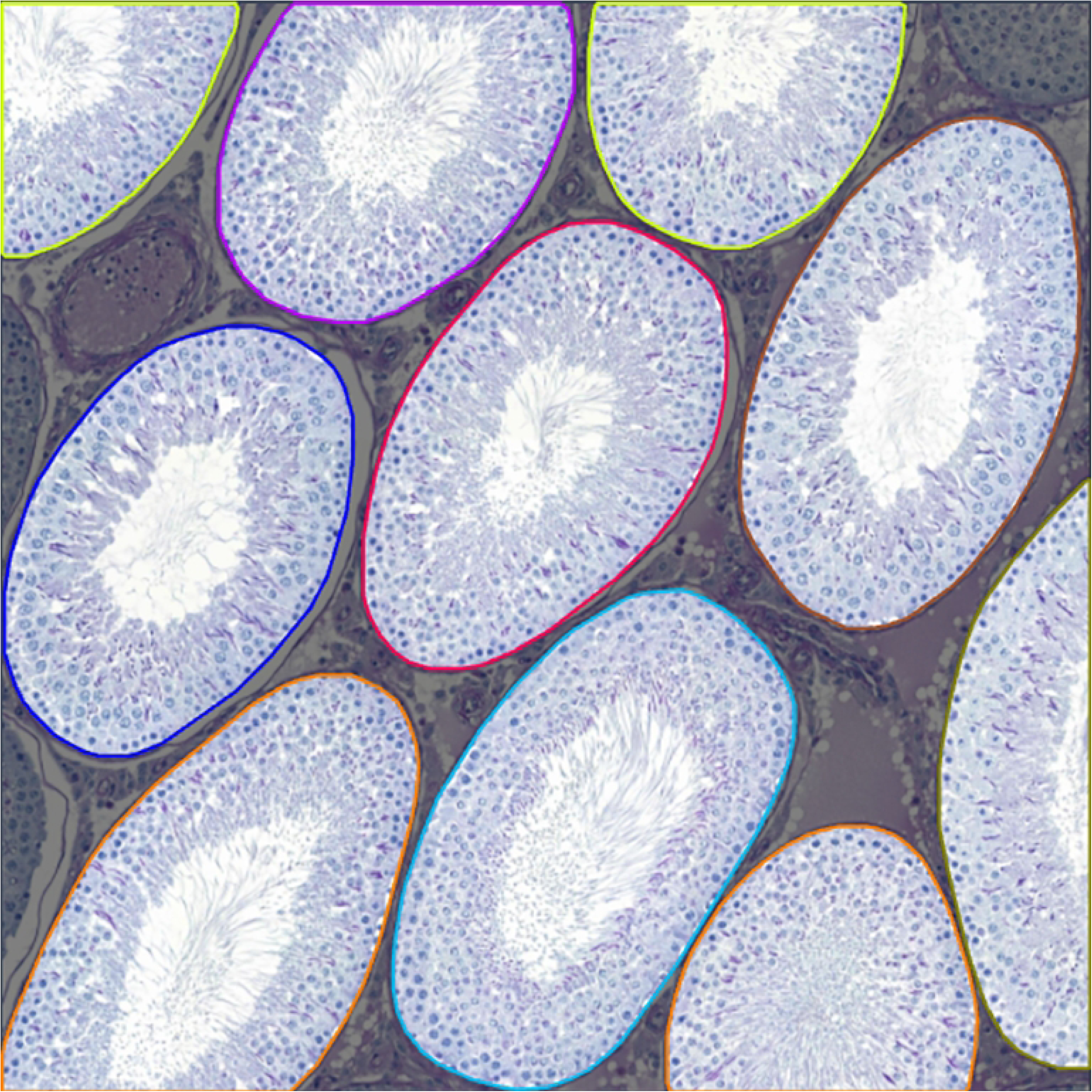
One example of class labeling on a tile image. Each class was distinguished by a specific color. The spermatogenic stages according to color were as follows —yellow green: stage I, red: stage II-III, purple: stage V, orange: stage VI, sky blue: stage VII, blue: stage XII, brown: stage XIII, khaki: stage XIV.

14 classes were identified, with spermatogenic stages II and III combined due to difficulty in microscopic differentiation. Atrophy was included as an additional class. The classification of stages was based on Russell’s standards (24). The criteria of spermatogenic stage are briefly illustrated in Table 1. Fig 2 shows the histology of spermatogenic stage.

**Fig 2.**
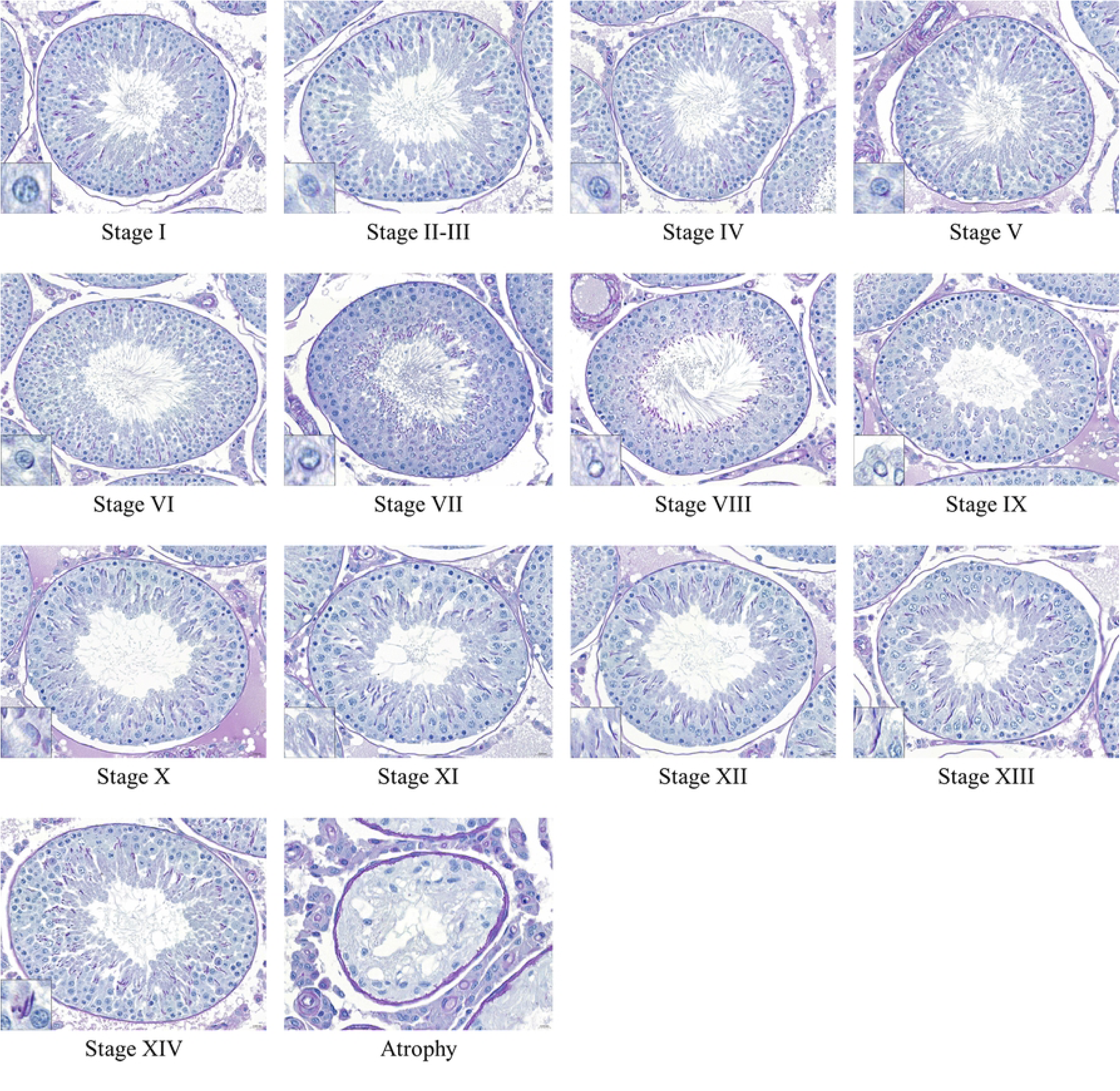
PAS-stained seminiferous tubules histology of spermatogenic stage. The 14 stages were classified based on the shape of acrosome and spermatid. Stages II and III were combined due to their indistinguishable appearance under microscopic observation. Atrophy was characterized by the depletion of germ cells and a reduction in the size of seminiferous tubules. The bottom-left box is a magnified view of acrosome and round spermatids.

**Table 1.**
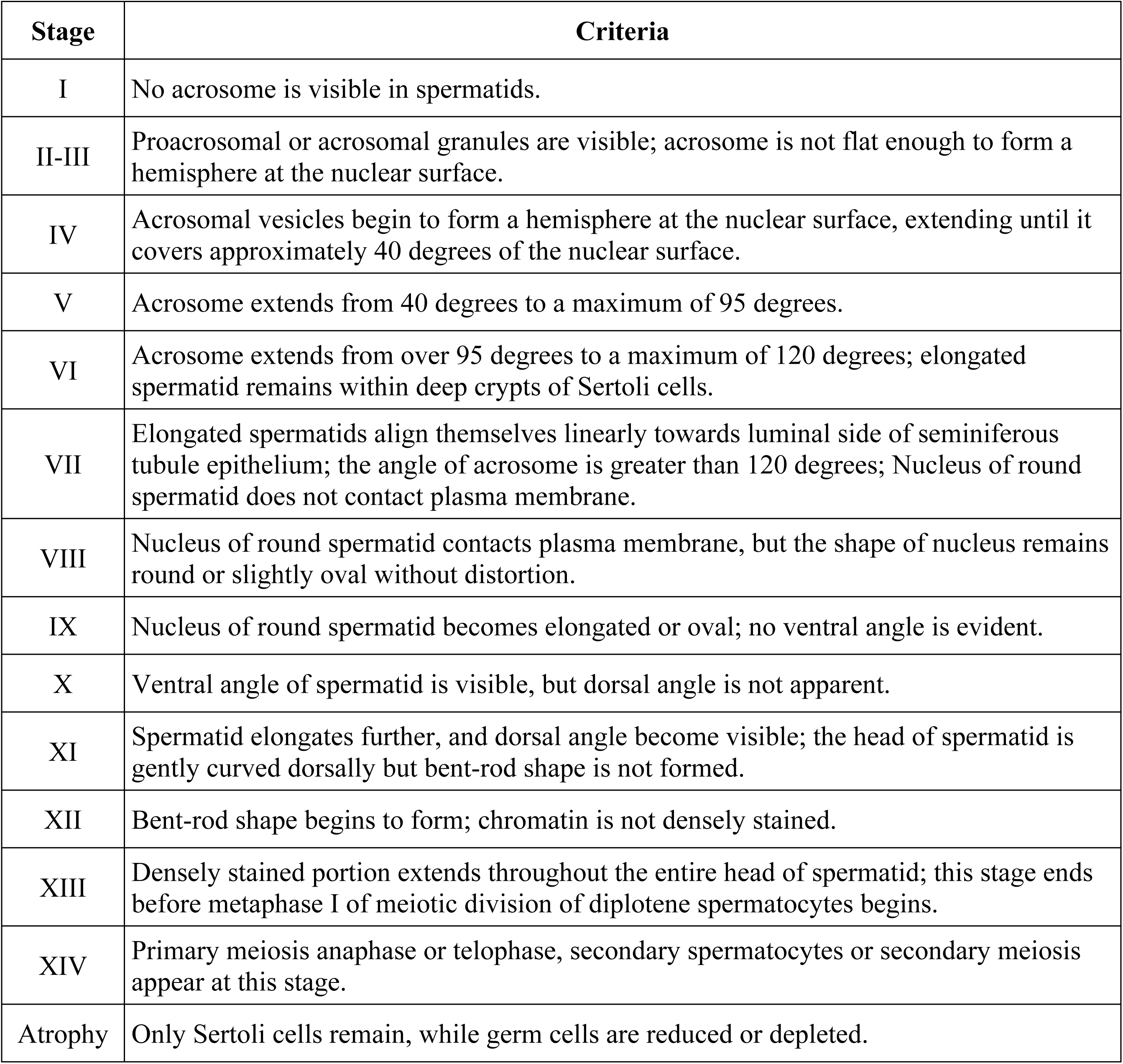
Criteria of spermatogenic stage. (^24^)

### Deep learning model training

#### Dataset preparation

The annotated 683 tile images were divided into train (70%, 478 images), valid (20%, 137 images), and test (10%, 68 images) datasets. Each image was downscaled to 2048×2048×3. Data augmentation techniques, including flipping (horizontal and vertical), 90-degree rotation (clockwise and counter-clockwise), cropping (0% minimum zoom and 25% maximum zoom), rotation (between −15° and +15°), shear (±10° horizontal and ±10° vertical), exposure (between −10% and +10%), and saturation (between −25% and +25%) adjustment, increased the training dataset to 1,434 images. One tile image contains multiple class objects as shown in Fig 1. Table 2 lists a total of 15,649 objects used for each dataset.

**Table 2.** Number of objects used in the models.

| Stage | Train | Valid | Test |
| --- | --- | --- | --- |
| I | 2458 | 198 | 118 |
| II-III | 1100 | 121 | 56 |
| IV | 624 | 73 | 41 |
| V | 649 | 77 | 27 |
| VI | 588 | 61 | 30 |
| VII | 2760 | 275 | 135 |
| VIII | 886 | 84 | 41 |
| IX | 439 | 45 | 24 |
| X | 402 | 33 | 17 |
| XI | 476 | 52 | 20 |
| XII | 1119 | 116 | 68 |
| XIII | 990 | 101 | 45 |
| XIV | 707 | 53 | 41 |
| Atrophy | 417 | 60 | 22 |
| Total | 13615 | 1349 | 685 |
|  | 15649 |  |  |

#### Comparison of model architecture

Faster R-CNN replaces inefficient Selective Search of Fast R-CNN with a Region Proposal Network (RPN), which directly generates region proposals from the shared feature maps of the backbone CNN. The RPN utilizes anchor boxes of various sizes and proportions to efficiently propose better regions. These proposals then go through Region of Interest (RoI) Pooling and fully connected layers for final classification and bounding box regression. Building on Faster R-CNN, Cascade R-CNN utilizes a multi-stage detection process to progressively refine region proposal. The first stage works like a typical Faster R-CNN detector, generating initial bounding box predictions. The second and third stages were trained using previous predicted boxes and a higher Intersection over Union (IoU) threshold than previous stages to reduce false positives. The simplified architecture of Faster R-CNN and Cascade R-CNN are shown in Fig 3.

**Fig 3.**
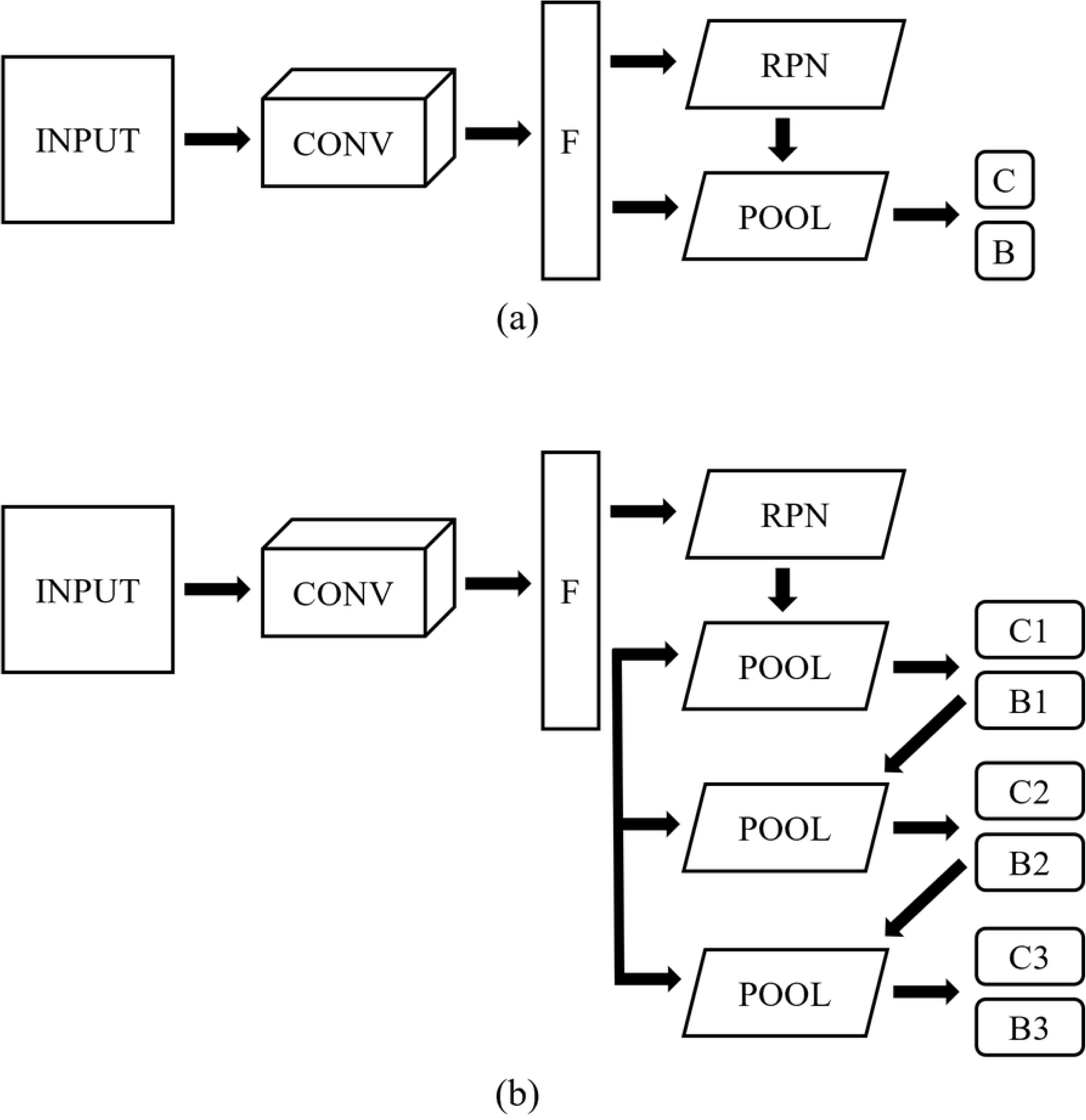
Faster R-CNN and Cascade R-CNN architectures. (a) Faster R-CNN architecture. The model directly and efficiently extracts region proposal from feature map through RPN. (b) Cascade R-CNN architecture. The model extracts region proposal similar to Faster-RCNN, but it utilizes a three stage detection process to increase precision. Abbreviations: CONV, convolutional layers; F, feature map; RPN, region proposal networks; POOL, RoI pooling; C, classification; B, bounding box regression.

#### Configuration settings

The scale of the training and testing images was set to 2048×2048. To manage memory, the batch size was set to 2 for training and 1 for validation and testing. The number of epochs was set to 12 or 24, with validation performed after each epoch. The learning rate was set to 0.02/10, with a linear learning rate of 0.001 applied for the first 500 iterations. Additionally, a learning rate decay was applied twice during training. Further details on the settings can be found in S1 File. Model training was conducted using an NVIDIA RTX A4500 GPU and the MMDetection toolbox (25).

### Model evaluation

#### Performance metrics

Precision and recall metrics were evaluated on the test dataset. Precision represents the proportion of true positives among the sum of true positives and false positives, while recall (sensitivity) quantifies true positives among the sum of true positives and false negatives.

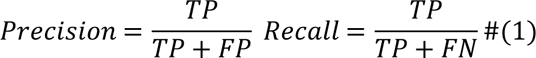

The Precision-Recall (PR) curve illustrates precision and recall variations according to confidence score thresholds. The Average Precision (AP), the area under the PR curve, evaluates model performance across confidence score thresholds. In multi-class scenarios, the mean Average Precision (mAP) is calculated. Similarly, mean Average Recall (mAR) represents the mean recall across IoU thresholds.

#### Whole slide image inference

To verify the practicality, inference was conducted on 6 WSIs not used in training or performance evaluation, including three normal (named normal A, B, and C) and three atrophied testes (severity of minimal, mild, severe). Atrophy grading was defined as follows: minimal (<10%), mild (10-30%), moderate (30-50%), and severe (>50%). Due to large WSI size, the SAHI library (26) was employed for memory-efficient inference. Inference was conducted using level 2 slide images, which provided the second-highest resolution among the 18 available levels. Due to computational memory constraints, level 2 was chosen as the optimal balance between resolution and processing capability. After WSI inference, the results of normal WSIs were compared with the evaluation of pathologist 1 who provided the ground truth.

#### Stage frequency and statistics

Stage frequencies were analyzed across the model, three pathologists (named pathologists 1, 2, and 3), and published literature by Hess et al. (27). Pathologist 1 was the ground truth provider who performed image annotation, and the other two pathologists 2 and 3 were independent pathologists unrelated to the ground truth. Statistical analysis of stage frequency was performed using IBM SPSS statistics software and the significance level was set at p<0.05. Multivariate Analysis of Variance (MANOVA) and Dunnett’s post-hoc test were used to compare stage-wise mean differences. The results of the model were set as a control group.

## Results

### Model performance metrics

The performance of the models was evaluated using the test dataset. Comparisons were made between two model types, Faster R-CNN and Cascade R-CNN, with two backbone options (ResNet-50 and ResNet-101) and two training durations (12 and 24 epochs). As shown in Table 3, the best-performing model was the Cascade R-CNN with a ResNet-50 backbone trained for 12 epochs. Increasing the backbone depth or extending training duration did not significantly improve performance. When the Intersection over Union (IoU) threshold range was set to 0.50:0.95, the best model achieved a mean Average Precision (mAP) of 0.869 and a mean Average Recall (mAR) of 0.977.

**Table 3.** Performance results of the models.

| Model type | Backbone | Epochs | mAP<sup>a</sup> | mAR<sup>a</sup> |
| --- | --- | --- | --- | --- |
| Faster R-CNN | ResNet-50 | 12 | 0.824 | 0.947 |
| Faster R-CNN | ResNet-101 | 12 | 0.807 | 0.959 |
| Faster R-CNN | ResNet-101 | 24 | 0.836 | 0.963 |
| Cascade R-CNN | ResNet-50 | 12 | 0.869 | 0.977 |
| Cascade R-CNN | ResNet-101 | 12 | 0.867 | 0.977 |
| Cascade R-CNN | ResNet-101 | 24 | 0.857 | 0.960 |

Abbreviations: mAP, mean Average Precision; mAR, mean Average Recall
<sup>a</sup>IoU=0.50:0.95

The result of stage-wise AP is represented in Fig 4. The AP of stages II–III, V, and XI was relatively lower. In case of atrophy detection, the AP tended to be high, exceeding 0.9, especially when using the Cascade R-CNN models.

**Fig 4.**
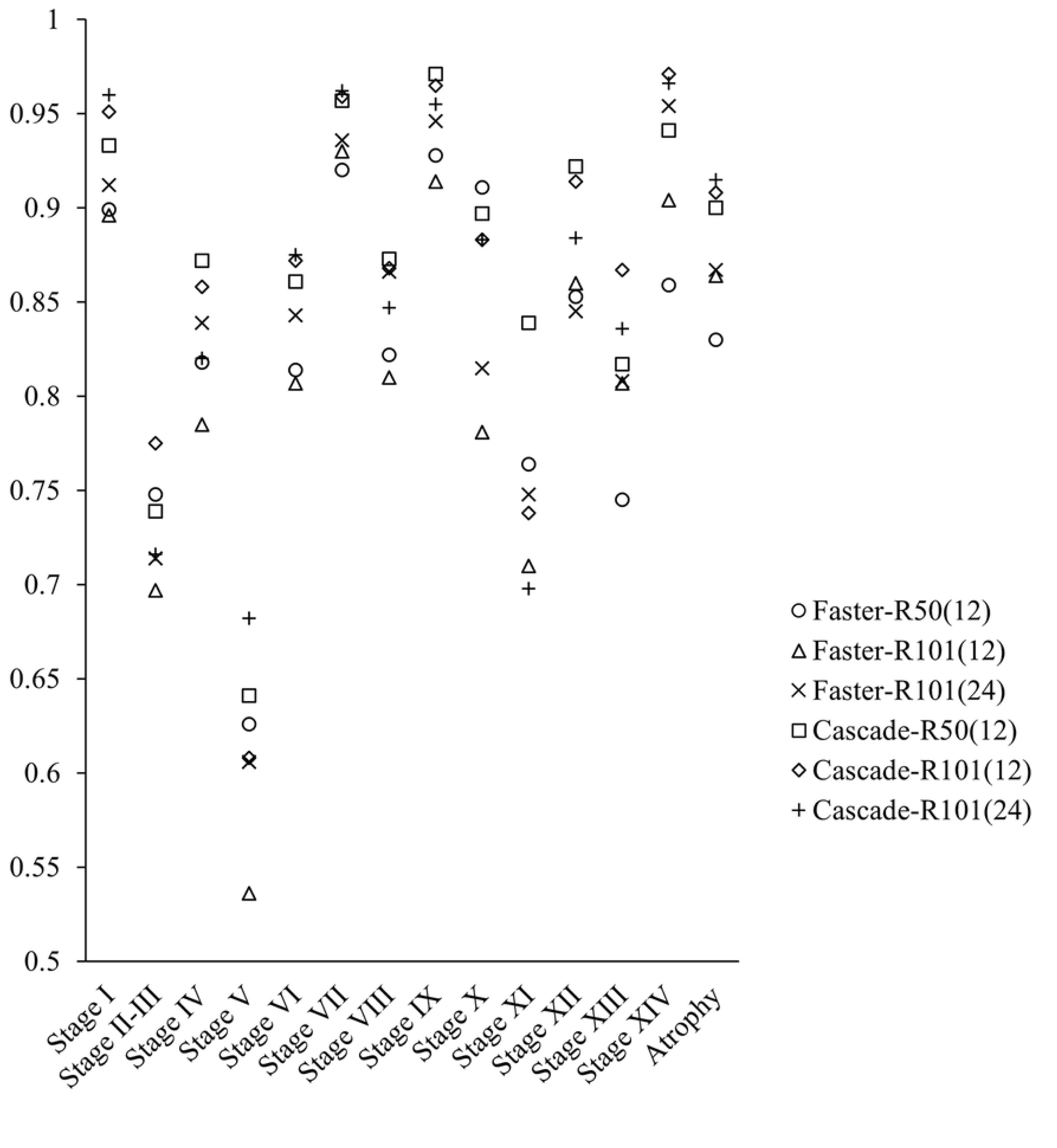
A dot plot showing stage-wise AP results of six models. The results of the Cascade R-CNN were generally higher than those of the Faster R-CNN. Legend: R is an abbreviation for ResNet, and

the numbers in parentheses represent epochs.

### Whole slide image inference for spermatogenic staging

The best-performing model (Cascade R-CNN with a ResNet-50 backbone) was used for WSI inference. The average inference time for three normal testicular WSIs was 211 seconds. Detailed inference time is presented in Table 4.

**Table 4.** WSI inference times for three normal testicular WSIs.

| WSI | Time (seconds) |
| --- | --- |
| Normal A | 216.01 |
| Normal B | 231.38 |
| Normal C | 185.80 |
| **Average** | 211.06 |

As shown in Fig 5, S1 Fig, and S2 Fig, the model successfully detected seminiferous tubules in normal testis WSIs. However, it occasionally produced duplicate inferences for longitudinally sectioned tubules. Duplicate bounding boxes accounted for an average of 1.92% of detections, while undetected tubules accounted for an average of 1.37%. Detailed proportion and count of duplicate and undetected boxes are shown in Table 5. Also, the model’s results were compared to pathologist 1’s assessments, who provided the ground truth, using a confusion matrix as shown in Fig 6. Detailed values for three slides are presented in S2 Table. Atrophied tubules near the rete testis were considered normal, so cases of atrophy detected in normal testes were excluded from the bar graph and confusion matrix results.

**Fig 5.**
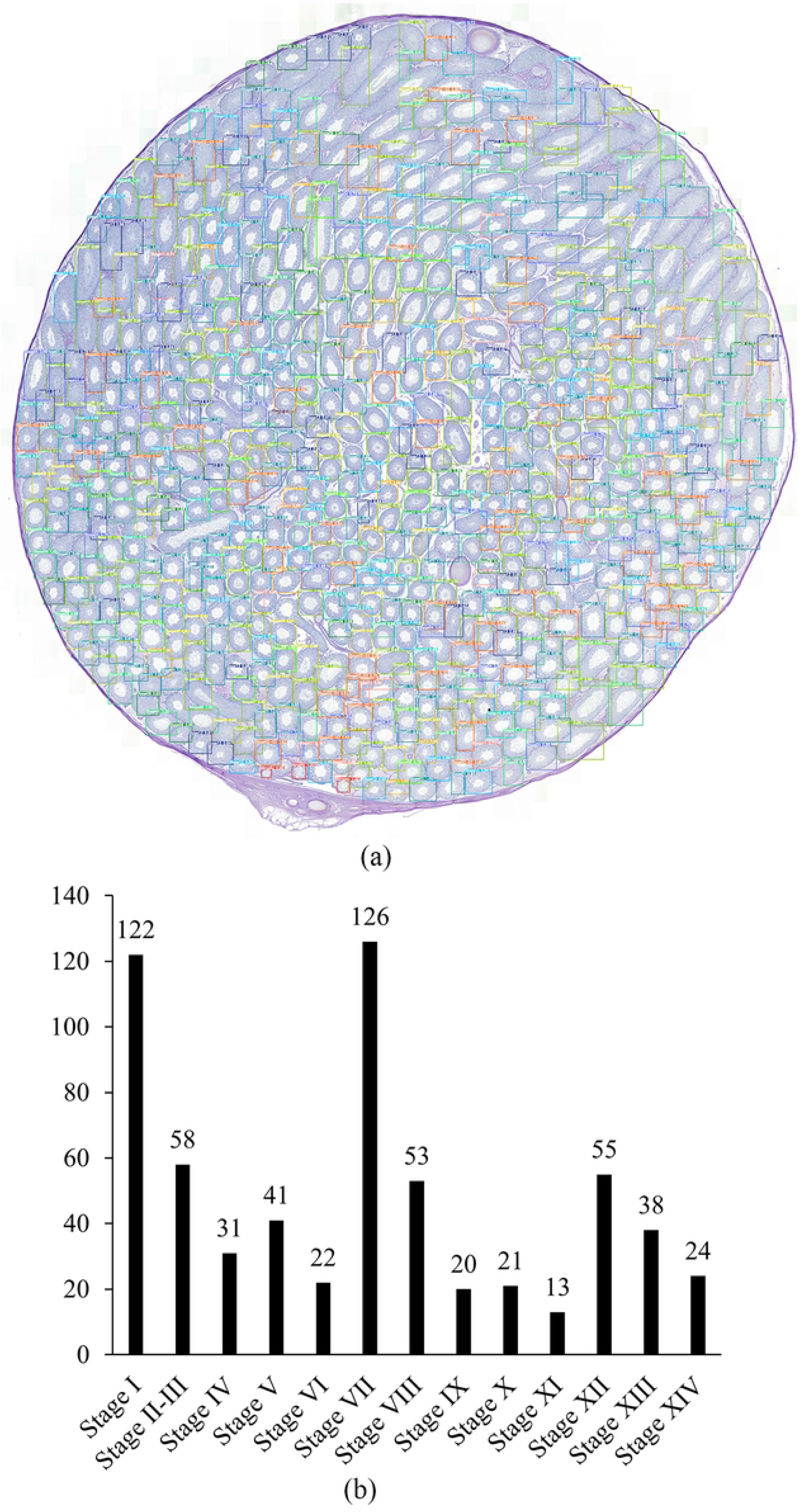
One of the results of normal testicular WSI inference (normal. **A)**. (a) WSI with inferred bounding boxes in different colors for each class. (b) bar graph showing the number of each class based on (a).

**Fig 6.**
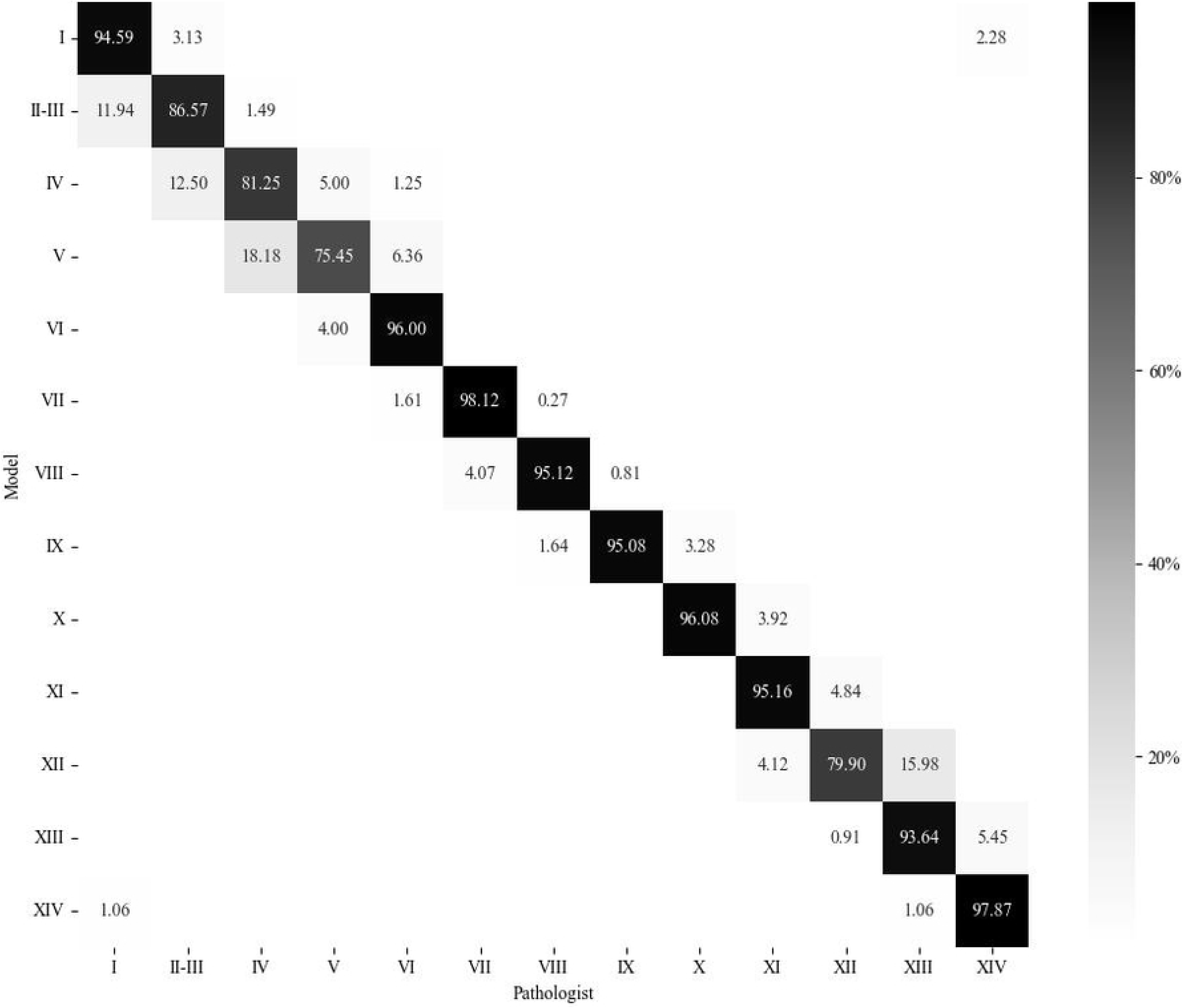
Confusion matrix comparing results of the model and the pathologist 1 in three normal testicular WSIs. The percentages represent precision (the proportion of true positive among the sum of true positive and false positive). Most stages demonstrated high accuracy, exceeding 90%, and no significant deviations from the ground truth were observed in the model’s output.

**Table 5.** Proportion and count of duplicate bounding boxes and undetected tubules in three normal testicular WSI inference results.

| WSI (total count) | Duplicate (count) | Undetected (count) |
| --- | --- | --- |
| Normal A (624) | 2.40% (15) | 1.44% (9) |
| Normal B (597) | 1.34% (8) | 1.50% (9) |
| Normal C (597) | 2.01% (12) | 1.17% (7) |
| **Average** | 1.92% | 1.37% |

Comparison of stage frequencies across the model, three pathologists, and the literature by Hess et al. is depicted in Fig 7. Detailed values of stage frequencies are presented in S3 Table. When compared the mean differences using statistical analysis, significant differences were found in the frequencies of stages I, VI, and XII between the model and the literature. However, no significant differences were observed between the model and the pathologists. Detailed values of statistics are presented in S4 Table.

**Fig 7.**
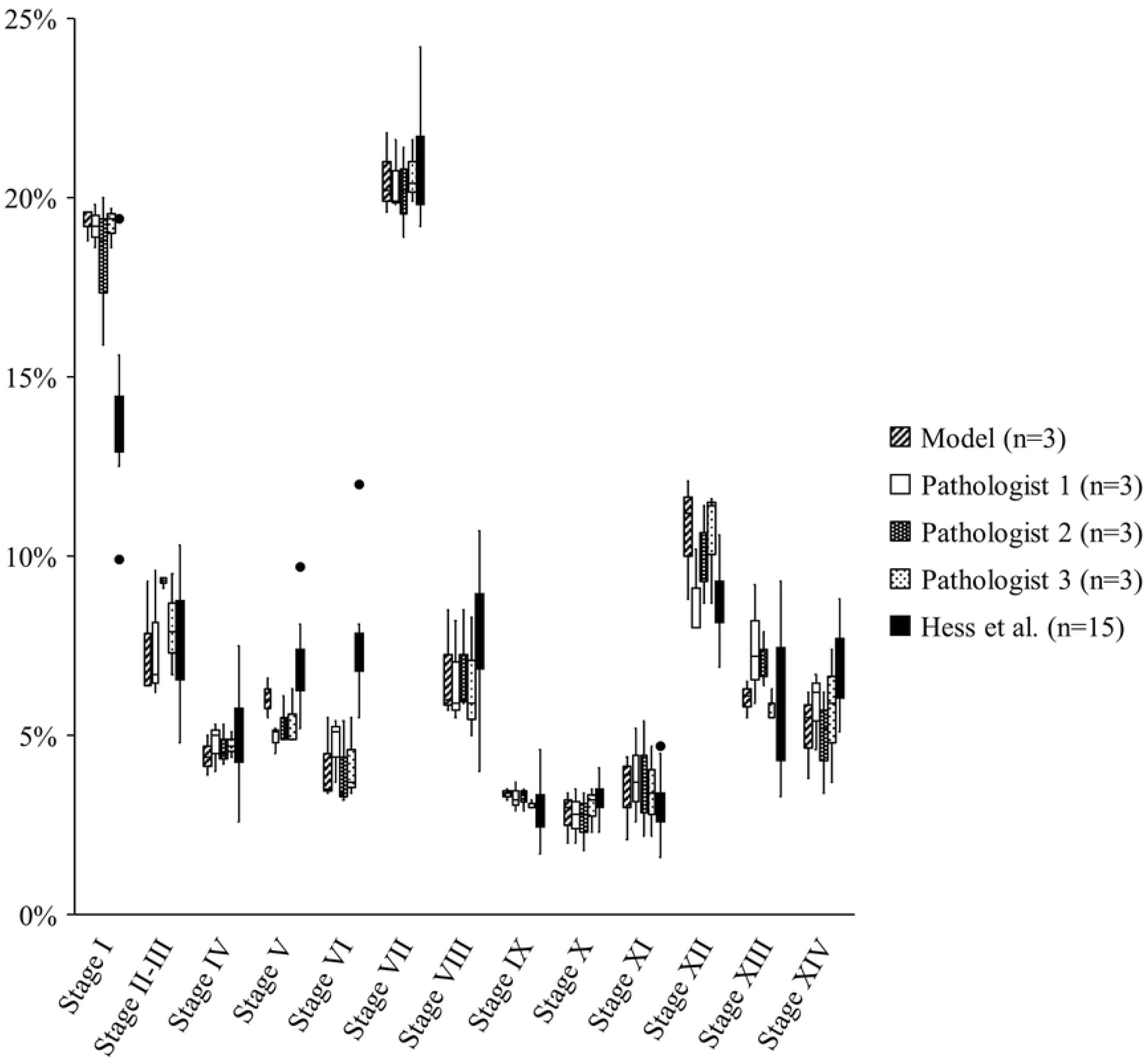
A box-and-whisker plot showing stage frequencies of the model, three pathologists, and the literature. The model and pathologists 1, 2, and 3 showed similar results, whereas differences were observed between the model and Hess et al.’s results in stages I, VI, and XII. This study evaluated spermatogenic stages in three testes, while the research by Hess et al. (1990) assessed stages in 15 testes. Pathologist 1 served as the ground truth, while pathologists 2 and 3 were independent pathology experts not associated with the ground truth.

### Whole slide image inference for atrophy detection

WSI inference was conducted for three atrophy severity grades—minimal, mild, and severe—to enable quantitative analysis. As shown in Fig 8-10, the model quite accurately detected atrophied seminiferous tubules across all severity levels. In severely atrophied testicular WSI, where all seminiferous tubules exhibited atrophy, only two misclassifications were observed.

**Figure 8.**
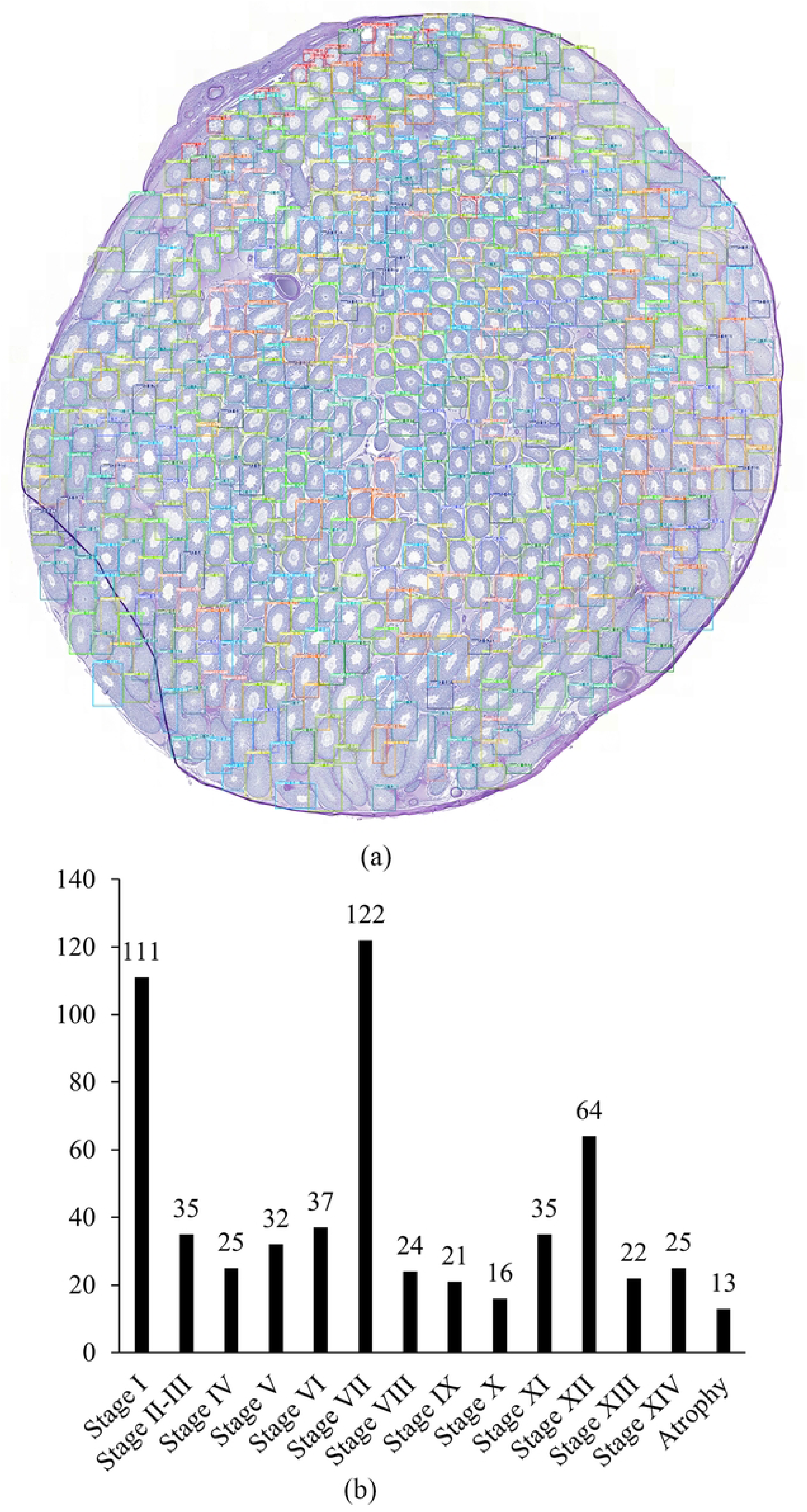
Minimal atrophied testicular WSI inference result. (a) WSI with inferred bounding boxes in different colors for each class. (b) bar graph showing the number of each class based on (a).

**Figure 9.**
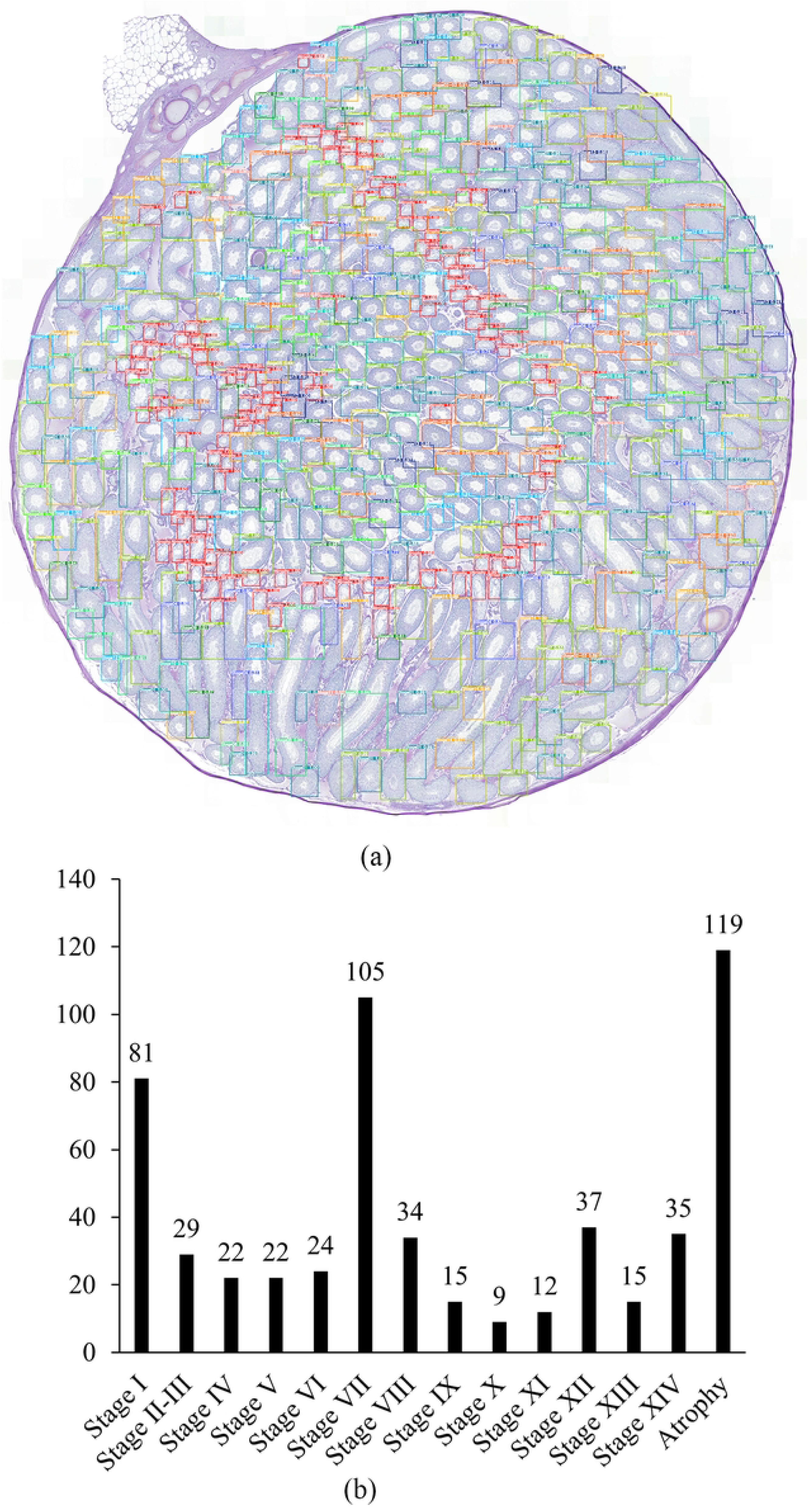
Mild atrophied testicular WSI inference result. (a) WSI with inferred bounding boxes in different colors for each class. (b) bar graph showing the number of each class based on (a).

**Figure 10.**
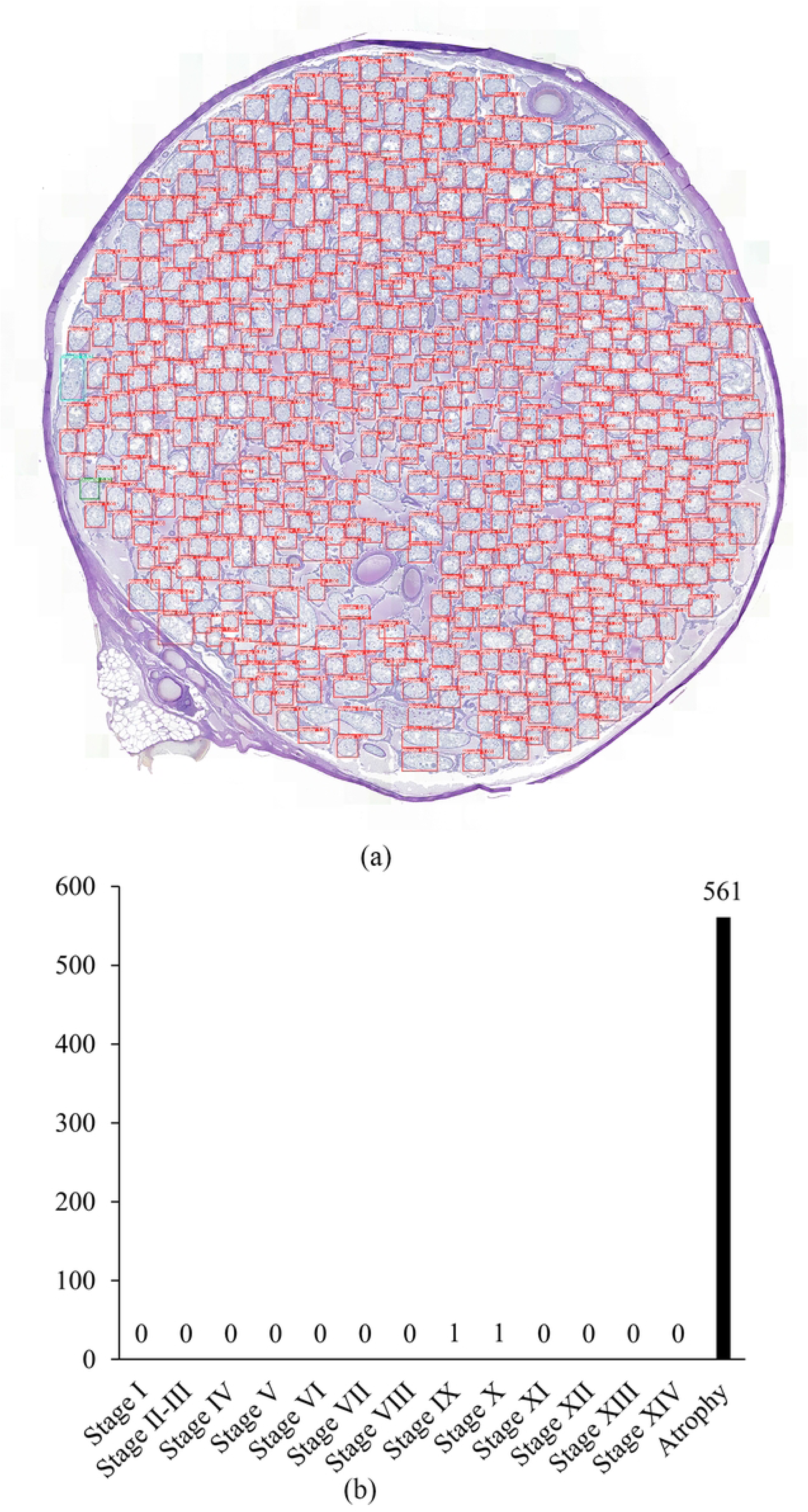
Severe atrophied testicular WSI inference result. (a) WSI with inferred bounding boxes in different colors for each class. (b) bar graph showing the number of each class based on (a).

## Discussion

The model performance metrics demonstrated that the best-performing model achieved a mAP of 0.869 and a mAR of 0.977. Higher recall suggests a tendency toward false positives, indicating that while the model effectively detects seminiferous tubules, it sometimes partially detects tubules or misclassifies their stages. This trend was also evident in WSI inference. In stage-wise AP analysis, relatively lower precision was observed for stages II–III, V, and XI. Stages II–III and V closely resemble adjacent stages (IV and VI), differing primarily in acrosome shape, which may have contributed to classification challenges. For stage XI, the distinctive morphology of spermatids is notable; however, the short duration of this stage resulted in fewer training samples and the co-occurrence of spermatids at different maturation stages. This may have caused misclassification between stages X, XI, and XII. Conversely, atrophy detection exhibited high AP (>0.9), suggesting the model’s potential for abnormal tissue analysis.

WSI inference results for normal testes indicated that the model successfully detected most seminiferous tubules with a low error rate (duplicate detections: 1.92%, undetected tubules: 1.37%). This suggests that automated spermatogenic stage assessment is feasible and could improve efficiency. Moreover, comparison of model results and pathologists’ evaluations demonstrated strong agreement, with a mean confusion matrix accuracy of 91%. The accuracy was higher than the mAP values, likely due to differences in evaluation criteria. Unlike AP calculations that rely on IoU and confidence scores, the confusion matrix considers a classification correct if the assigned label is accurate, even when bounding box overlap is partial. Stage frequency analysis revealed that stages I and XII were more frequent, while stage VI was less frequent compared to the literature by Hess et al.. In the literature, fifteen rats aged approximately 13-16 weeks were used, whereas in this study, three rats aged 21 weeks were used. Therefore, further research with a larger sample size is needed to determine whether differences in stage frequency are influenced by the age of the rats, housing conditions, or other factors. Although frequency differences exist, deep learning models provide consistent classification criteria, which are valuable for toxicological assessments that focus on comparative rather than absolute evaluations. The absence of significant differences in frequency between the model and pathologists further supports its reliability.

In atrophied testis inferences, atrophy detection was highly accurate. In severely atrophied testis, only 2 seminiferous tubules were misclassified as stages IX and X. This may be due to the morphological similarity between atrophied tubules and stages IX and X, where only round spermatids are present and the lumen appears empty. However, the low misclassification rate suggests that the model does not significantly impact atrophy grading. Currently, atrophy grading relies on area-based assessment, which is somewhat subjective. The deep learning model provides a precise atrophied tubule count, allowing for more objective and quantitative grading.

## Conclusion

This study investigated the feasibility of using deep learning models for automated spermatogenic stage classification in rat testes. The object detection model demonstrated high accuracy and efficiency in WSI inference, providing results comparable to those of pathologists. In normal testicular tissues, the model reliably identified all 14 spermatogenic stages, aiding pathologists by offering an objective reference and improving workflow efficiency. It also facilitated comparative analysis across multiple samples with varying seminiferous tubule counts. Further research on the spermatogenic stage frequency is needed, and the efficiency of the model can aid in this process. In atrophied testicular tissues, the model effectively quantified atrophied tubules, supporting objective grading through quantitative analysis. Overall, the object detection model showed strong applicability to real-world tasks, significantly enhancing accuracy and efficiency in spermatogenic stage evaluation. By leveraging deep learning, pathologists can conduct rapid and precise assessments without excessive time investment. Furthermore, unlike conventional subjective methods, the model provides an objective standard for toxicological assessments. Automating routine tasks with deep learning allows pathologists to focus on more complex analyses, improving overall diagnostic efficiency. Future research expanding the application of deep learning models to other pathological conditions could further enhance their utility and impact in toxicopathological research.

## Acknowledgments

The authors thank Tae-Kyun Kim for advice on deep learning-based data analysis.

## Supporting information

**S1 Fig. Normal B WSI inference result.** (a) WSI with inferred bounding boxes in different colors for each class. (b) bar graph showing the number of each class based on (a).

**S2 Fig. Normal C WSI inference result.** (a) WSI with inferred bounding boxes in different colors for each class. (b) bar graph showing the number of each class based on (a).

**S1 Table. Stage-wise AP result of six models.** Abbreviations: AP, Average Precision, AR, Average Recall

**S2 Table. Confusion matrix between the model and pathologist 1 (ground truth).** (a) confusion matrix of normal A testicular WSI. (b) confusion matrix of normal B testicular WSI. (c) confusion matrix of normal C testicular WSI. (d) confusion matrix of sum of all three normal testicular WSIs.

**S3 Table. Stage frequency values for the evaluation results of the model, three pathologists, and Hess et al.** Spermatogenic stage frequencies were calculated from normal testicular tissues.

**S4 Table. Statistics of stage frequency.** Significance was assessed when the p value of the mean difference was less than 0.05.

**S1 File. System environment and configuration settings.**

## References

OECD. Test No. 421: Reproduction/Developmental Toxicity Screening Test2016.

OECD. Test No. 422: Combined Repeated Dose Toxicity Study with the Reproduction/Developmental Toxicity Screening Test2016.

Kim I, Kang K, Song Y, Kim T-J. Application of Artificial Intelligence in Pathology: Trends and Challenges. Diagnostics. 2022;12(11):2794.

Shafi S, Parwani VA. Artificial intelligence in diagnostic pathology. Diagnostic Pathology. 2023;18(1).

De Rosa L, L’Abbate S, Kusmic C, Faita F. Applications of Deep Learning Algorithms to Ultrasound Imaging Analysis in Preclinical Studies on In Vivo Animals. Life (Basel). 2023;13(8).

Küper A, Blanc-Durand P, Gafita A, Kersting D, Fendler WP, Seibold C, et al. Is There a Role of Artificial Intelligence in Preclinical Imaging? Seminars in Nuclear Medicine. 2023;53(5):687–93.

Creasy DM, Panchal ST, Garg R, Samanta P. Deep Learning-Based Spermatogenic Staging Assessment for Hematoxylin and Eosin-Stained Sections of Rat Testes. Toxicol Pathol. 2021;49(4):872–87.

Mehrvar S, Kambara T. Morphologic Features and Deep Learning-Based Analysis of Canine Spermatogenic Stages. Toxicol Pathol. 2022;50(6):736–53.

Xu J, Lu H, Li H, Yan C, Wang X, Zang M, et al. Computerized spermatogenesis staging (CSS) of mouse testis sections via quantitative histomorphological analysis. Med Image Anal. 2021;70:101835.

Mecklenburg L, Luetjens CM, Romeike A, Garg R, Samanta P, Mohanty A, et al. Deep Learning–Based Spermatogenic Staging in Tissue Sections of Cynomolgus Macaque Testes. Toxicologic Pathology. 2024;52(1):4–12.

Girshick R, Donahue J, Darrell T, Malik J, editors. Rich feature hierarchies for accurate object detection and semantic segmentation. Proceedings of the IEEE conference on computer vision and pattern recognition; 2014.

Ren S, He K, Girshick R, Sun J. Faster R-CNN: Towards Real-Time Object Detection with Region Proposal Networks. IEEE Trans Pattern Anal Mach Intell. 2017;39(6):1137–49.

Cai Z, Vasconcelos N. Cascade R-CNN: High quality object detection and instance segmentation. IEEE transactions on pattern analysis and machine intelligence. 2019;43(5):1483–98.

Foley GL. Overview of male reproductive pathology. 2001;29(1):49–63.

Leblond CP, Clermont Y. Definition of the stages of the cycle of the seminiferous epithelium in the rat. Ann N Y Acad Sci. 1952;55(4):548–73.

Clermont Y, Perey B. The stages of the cycle of the seminiferous epithelium of the rat: practical definitions in PA-Schiff-hematoxylin and hematoxylin-eosin stained sections. Rev Can Biol. 1957;16(4):451–62.

Narayana K, Prashanthi N, Nayanatara A, Bairy LK, D’Souza UJ. An organophosphate insecticide methyl parathion (o-o-dimethyl o-4-nitrophenyl phosphorothioate) induces cytotoxic damage and tubular atrophy in the testis despite elevated testosterone level in the rat. J Toxicol Sci. 2006;31(3):177–89.

Oishi S. Testicular atrophy induced by di(2-ethylhexyl)phthalate: changes in histology, cell specific enzyme activities and zinc concentrations in rat testis. Arch Toxicol. 1986;59(4):290–5.

Oliveira CA, Carnes K, França LR, Hess RA. Infertility and testicular atrophy in the antiestrogen-treated adult male rat. Biol Reprod. 2001;65(3):913–20.

Creasy D, Bube A, de Rijk E, Kandori H, Kuwahara M, Masson R, et al. Proliferative and nonproliferative lesions of the rat and mouse male reproductive system. Toxicol Pathol. 2012;40(6 Suppl):40S–121S.

O’Donnell L, McLachlan R, Wreford N, Robertson D. Testosterone promotes the conversion of round spermatids between stages VII and VIII of the rat spermatogenic cycle. Endocrinology. 1994;135(6):2608–14.

Kerr J, Millar M, Maddocks S, Sharpe R. Stage-dependent changes in spermatogenesis and Sertoli cells in relation to the onset of spermatogenic failure following withdrawal of testosterone. The Anatomical Record. 1993;235(4):547–59.

Goode A, Gilbert B, Harkes J, Jukic D, Satyanarayanan M. OpenSlide: A vendor-neutral software foundation for digital pathology. J Pathol Inform. 2013;4:27.

L. D. Russell RAE, A. P. S. Hikim and E. D. Clegg. Histological and Histopathological Evaluation of The Testis: Cache River Press; 1990.

Chen K, Wang J, Pang J, Cao Y, Xiong Y, Li X, et al. MMDetection: Open mmlab detection toolbox and benchmark. arXiv preprint arXiv:190607155. 2019.

Akyon FC, Cengiz C, Altinuc SO, Cavusoglu D, Sahin K, Eryuksel O. SAHI: A lightweight vision library for performing large scale object detection and instance segmentation. November; 2021.

Hess RA, Schaeffer DJ, Eroschenko VP, Keen JE. Frequency of the stages in the cycle of the seminiferous epithelium in the rat. Biology of reproduction. 1990;43(3):517–24.

